# Zelda potentiates transcription factor binding to zygotic enhancers by increasing local chromatin accessibility during early *Drosophila melanogaster* embryogenesis

**DOI:** 10.1101/380857

**Authors:** Xiao-Yong Li, Michael B. Eisen

## Abstract

The maternally deposited transcription factor Zelda binds to and is required for the activation of a large number of genes in early *Drosophila* development, and has been suggested to act as a pioneer factor. In this study, we investigated the temporal dynamics of Zelda binding along with the maternal patterning factors Dorsal and Caudal during early embryogenesis. We found in regions bound by Zelda and either Dorsal or Caudal, Zelda binding was detected, and reached maximum levels, earlier than Caudal and Dorsal, providing support of its role as a pioneer factor. We found that Dorsal and Caudal binding correlated strongly with Zelda binding at mitotic cycle 12, suggesting that Zelda is important for early binding by these factors and early onset of their target gene expression. At the same time, we show that among Dorsal target enhancers, the dorsal and ventral ectoderm enhancers are much more strongly associated with Zelda than mesoderm enhancers, revealing an additional function of Zelda in coordinating spatial activity of enhancers. We have also investigated the role of Zelda on chromatin structure. We found that in early embryos, before Dorsal and Caudal are bound at significant levels, Zelda binding is associated with histone acetylation and local histone depletion. These chromatin associated changes accompanied with increased local chromatin accessibility were also detected around Zelda peaks in coding sequences that do not appear to play a role in subsequent transcription factor binding. These findings suggest that Zelda binding itself can lead to chromatin structural changes. Finally, we found that Zelda motifs, both bound and unbound, tend to be associated with positioned nucleosomes, which we suggest may be important for the regulatory specificity of enhancers.

## Introduction

Enhancers that control spatial and temporal patterns of gene expression during animal development are typically bound by multiple transcription factors, which allow for the integration of intracellular and extracellular regulatory signals (Lelli et al., 2012; Spitz and Furlong, 2012). Transcription factors mediate enhancer function in several ways, with some directly regulating the recruitment of RNA polymerase and others influencing the ability of other factors to bind to the enhancer (Hashimoto et al., 2014). In particular, a group of transcription factors known as pioneer factors (Zaret and Carroll, 2011; Zaret and Mango, 2016) have been shown to regulate enhancer activity by binding to loci that are in a “closed” chromatin state - where most transcription factors are unable to bind - and subsequently, rendering them accessible to binding.

We have previously shown that the maternally deposited *Drosophila melanogaster* transcription factor Zelda (Zld) (Liang et al., 2008) exhibits pioneerlike activity at enhancers active during early embryogenesis (Harrison et al., 2011; Li et al., 2014). Zld binding sites are the most highly enriched sequences in early embryonic enhancers(Bosch, 2006; Li et al., 2008). Zld binds to a large fraction of enhancers active in the early embryo prior to their involvement in transcriptional regulation (Harrison et al., 2011) and early Zld binding is associated with reduced nucleosome density and increased histone acetylation (Li et al., 2014). Finally, Zld is also required for chromatin accessibility at zygotic enhancers (Foo et al., 2014; Li et al., 2014; Schulz et al., 2015; Sun et al., 2015).

While these observations strongly suggest that Zld is a pioneer factor, whether Zld binding precedes the binding of other early acting transcription factors has yet to be directly demonstrated. Mechanistically, Zld’s precise role in modulating chromatin structure is unknown. To address these questions, we used chromatin-immunoprecipitation coupled with high-throughput DNA sequencing (ChlP-seq) to analyze the temporal dynamics of Zld binding and the binding of two major, maternal factors that initiate the zygotic gene activation in early embryogenesis - Caudal(Cad) and Dorsal (Dl). Cad forms a broad posterior gradient in early embryo (Macdonald and Struhl, 2004)) and is important for patterning the embryo trunk along the anterior - posterior axis, while Dl forms a ventral to dorsal morphogen gradient, and drives sequential patterning along the dorsal-ventral axis(Stathopoulos and Levine, 2004).

## Results

### Zld, Cad, and Dl display different overall binding dynamics during the maternal to zygotic transition

To analyze the temporal dynamics of genome wide binding by Zld, Dl, and Cad during early fly embryogenesis, we carried out ChlP-seq experiments as previously described (Li et al., 2014) (Fig. 1A). Briefly, we hand-sorted formaldehyde-fixed embryos to obtain embryos of defined stages at approximately mitotic cycles 8, 12, 14a, and 14c. We isolated chromatin from these embryos and performed chromatin immunoprecipitation using antibodies against these factors. Prior to immunoprecipitation, we added a defined amount of chromatin from *D. pseudoobscura* stage 4/5 embryos to serve as normalization control.

**Fig.1.**
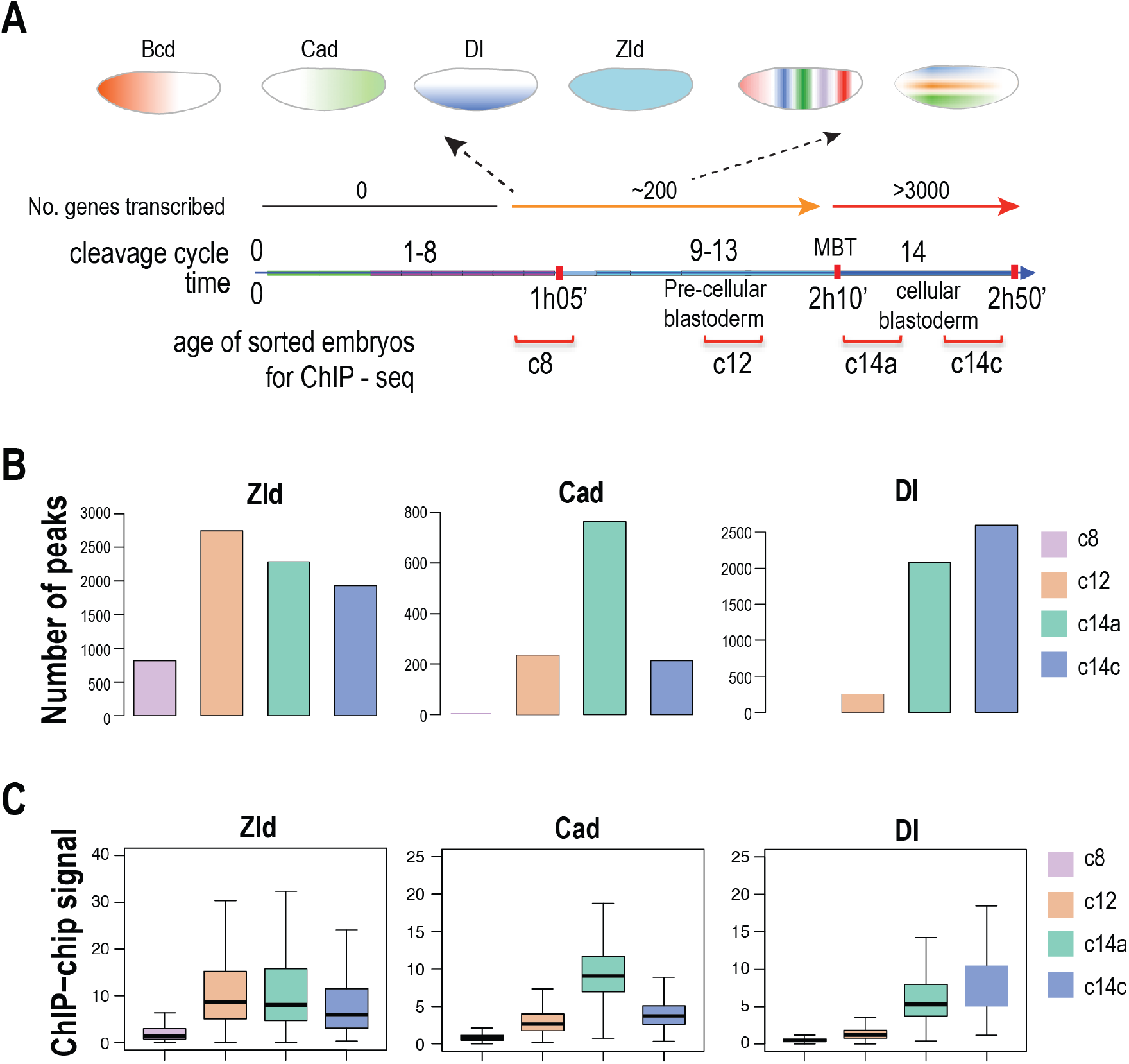
Overall trend of transcription factor binding during maternal to zygotic transition. A. Schematic for the major events during early embryo development including: rapid nuclear division in the 1^st^ one hour, the formation of the anterior - posterior (Bcd, Cad) and dorsal - ventral gradient (Dl) of maternal patterning factors, the onset of the minor wave of zygotic gene expression around mitotic cycle 8, and the start of cellularization at mid blastula transition, which marks the end of maternal to zygotic transition and the start of full range of transcription. The distributions of the ages of the sorted embryos (similar to previously described(Li et al., 2014)) used for chromatin preparation for ChIP-seq are as shown. B. The numbers of peaks identified in the ChIP-seq experiments for each factor using the peak calling program MACS (Zhang et al., 2008) are shown. C. The box plots show the distribution of the ChIP-seq signal around the peaks of each factor at each developmental stage of the embryo and the trend over time.

After aligning sequencing reads to the genome, we determined enriched peaks using the program MACS (Zhang et al., 2008), generated a consolidated peak list for each factor by merging peaks called from all stages, and determined the ChIP signal for each peak at each developmental stage. The signal from each stage was normalized based on the number of reads that aligned to the *D. pseudoobscura* genome, as described previously (Li et al., 2014).

As shown in Fig.1B and 1C, the three factors displayed distinct temporal dynamics. Zld binding was detected early, as expected from our previous study (Harrison et al., 2011), with hundreds of peaks detected at cycle 8, and reached its maximum at cycle 12. In contrast, there was little or no binding of Cad or Dl at cycle 8. Cad and Dl binding became detectable starting at cycle 12, and reached maximum at cycle 14a and cycle cycle 14c, respectively. The overall binding dynamics of these factors matched their known overall protein level dynamics between cycle 8 to cycle 14. Specifically, previous studies showed that even though the Dl gradient forms relatively early, it progresses slowly, reaches its maximum at the start of nuclear cycle 14, and remains stable during most of the cycle (Kanodia et al., 2009). The temporal spatial distribution of Cad is more complex and dynamic: it forms a broad gradient along the posterior – anterior axis starting as early as cycle 8; the gradient increases in intensity and reaches maximum by early cycle 14; in cycle 14, the gradient retracts gradually from the anterior, ending with a narrow posterior stripe remaining in late cycle 14 (Macdonald and Struhl, 2004).

### In early embryos, Zld binding precedes, and potentiates, Dorsal binding

We first investigated the potential impact of Zld binding on Dl, by comparing their dyanmics at sites with different temporal patterns of Dl binding. The 246 Dl peaks detected at cycle 12 were generally associated with strong Zld binding, in contrast to the approximately 2,000 Dl peaks detected at cycle 14a and cycle 14c (Fig.2A,B). Zld binding at sites overlapping many Dl peaks was detected at cycle 8, and on average achieved maximum binding at cycle 12, when Dl binding in general is still weak, as shown in Fig. 2B,D and as exemplified by the *sog* shadow enhancer (Hong et al., 2008) and *zen* dorsal ectoderm enhancer (Markstein, 2004) (Fig. 2C), consistent with a role for Zld as a pioneer factor potentiating the subsequent binding of these factors.

**Fig.2.**
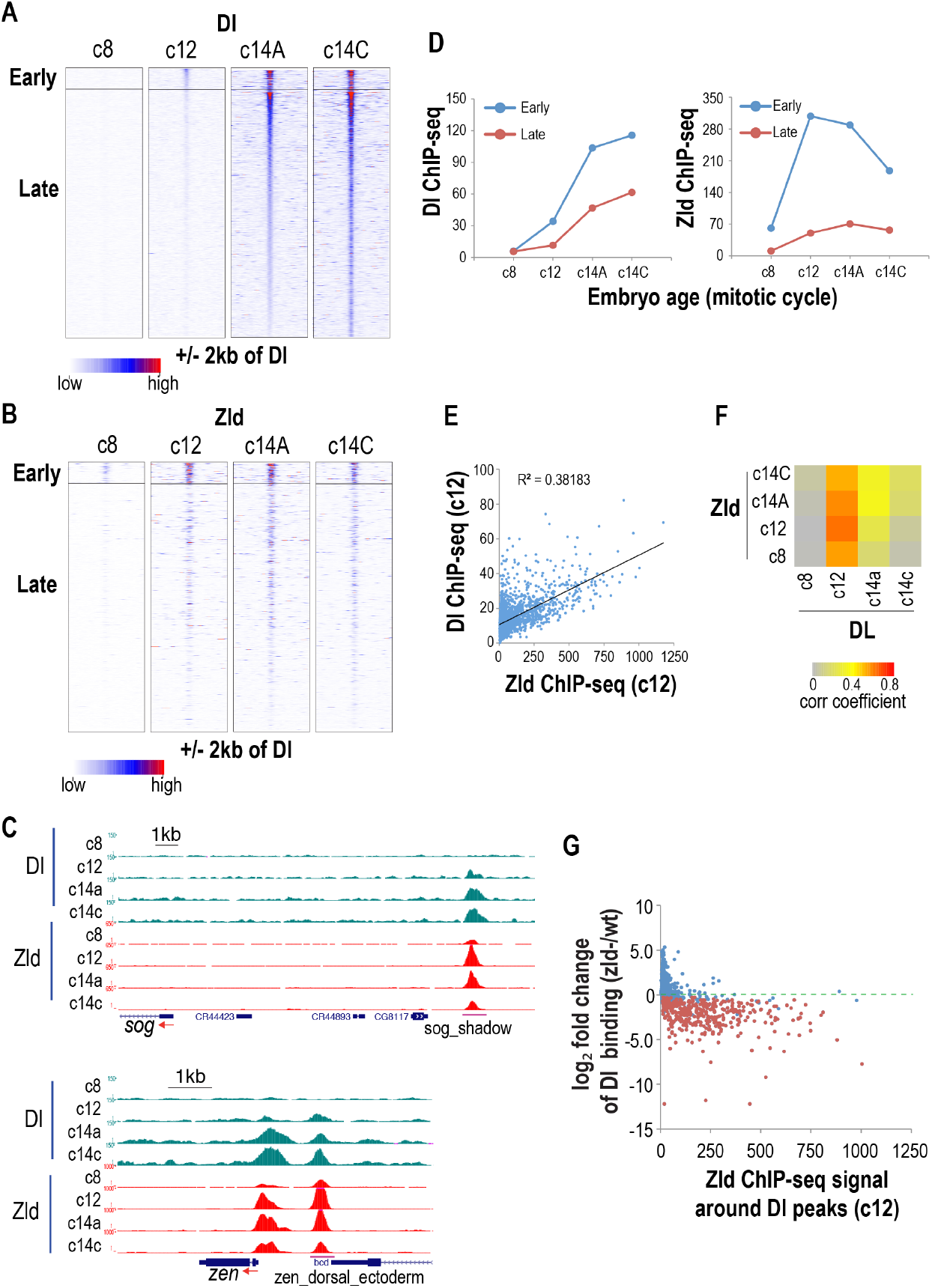
Relationship between the dynamics of Dl and Zld binding dynamics. A. Heatmap of ChIP-seq signals of the Dl peaks at the different stages of early embryo development. Early binding peaks, which were detected in cycle 12, were sorted according to ChIP-seq signal strength at cycle 12, and the late binding peaks were sorted according to ChIP-seq signal at cycle 14c. B. Heatmap of ChIP-seq signals of Zld around the Dl peaks. The peaks are grouped and sorted as in A. C. Genome browser snapshot of genomic regions around *sog* shadow/NEE enhancer (Crocker et al., 2008; Hong et al., 2008) and the *zen* dorsal ectoderm enhancer(Markstein, 2004), showing the ChIP-seq profiles of Zld and Dl at the different stages of early embryo development. D. Trends of the average ChIP-seq signal for Dl and Zld around the early and late groups of Dl peaks. E. Scatter plot showing correlation between the ChIP-seq signals of Dl and Zld around the Dl peaks at cycle 12. F. Heatmap showing the Pearson correlation coefficients between ChIP-seq signal of Dl and Zld around Dl peaks at each stage of early embryo development. G. fold change (in log_2_ scale) of Dl ChIP-seq signal in *zld*- mutant relative to wild type embryos (Sun et al., 2015) is plotted against Zld ChIP-seq signal detected around the Dl peaks at cycle 12, showing the relationship between level of Zld binding and the dependence of Dl binding on Zld at each Dl binding site.

We also observed a strong correlation between the levels of Dl and Zld binding at common targets in cycle 12, and, more weakly, at cycle 14a and cycle 14c (Fig. 2E and F), suggesting a direct role for Zld in mediating Dl binding. To investiage this further, we reanalyzed previously published data on Dl binding in *zld*- embryos (Sun et al., 2015), and found that the majority of the Dl peaks associated with Zld binding, particularly those at cycle 8 and 12 showed a strong decrease in binding in *zld*- mutant embryos (Fig. 2G, Fig. S1). Together with the finding that Zld binds to its targets in the absence of Dl at cycle 8, this demonstrates that there is likely a causal relationship between Zld binding and Dl binding in early embryos. In addition, the correlation between Dl binding and Zld binding suggests that Zld may play a role in controlling Dl activation levels, and therefore in regulating the temporal dynamics of dorsal target gene expression during zygotic genome activation, as was previously shown by (Sandler and Stathopoulos, 2016).

### Dependence of Dl binding on Zld correlates with spatial patterns of its target genes and enhancers

Dl forms a dorsal-ventral gradient, and drives expression of target genes with distinct dorsal-ventral patterns. Zld is speculated to be important for boosting the activity of Dl in the lateral and dorsal part of the embryo where the nuclear Dl concentration is low (Foo et al., 2014). To investigate this possibility, we curated a list of 83 genes (Table S1) that represent the three classic classes of dorsal-ventral genes (Stathopoulos and Levine, 2004) with distinct expression patterns in the presumptive mesoderm, ventral ectoderm, and dorsal ectoderm/dorsal amnioserosa, respectively, based on published studies (Leptin, 1991; Sandmann et al., 2007; Stathopoulos et al., 2002) and expression patterns in the BDGP *in situ* database (Hammonds et al., 2013; Tomancak et al., 2007).

Dl peaks associated with genes in this curated list were often associated with Zld binding (Fig. 3A), with Zld binding already present at cycle 8 when Dl binding was not yet detectable. Zld binding was observed over Dl peaks of all three classes of genes. Even so, by dividing the Dl peaks into three equal-sized groups based on levels of Zld binding at cycle 12, we found that while the majority of Dl peaks associated with the dorsal (dorsal ectoderm/dorsal amnioserosa genes, and large portion of those associated with the ventral ectoderm genes, are associated with strong or modest levels of Zld binding, relative few associated with the mesoderm genes are(Fig. 3B). This suggests that dorsal and lateral patterned genes have a stronger dependence on Zld than genes expressed in ventral (mesoderm) part of the embryo.

**Fig.3.**
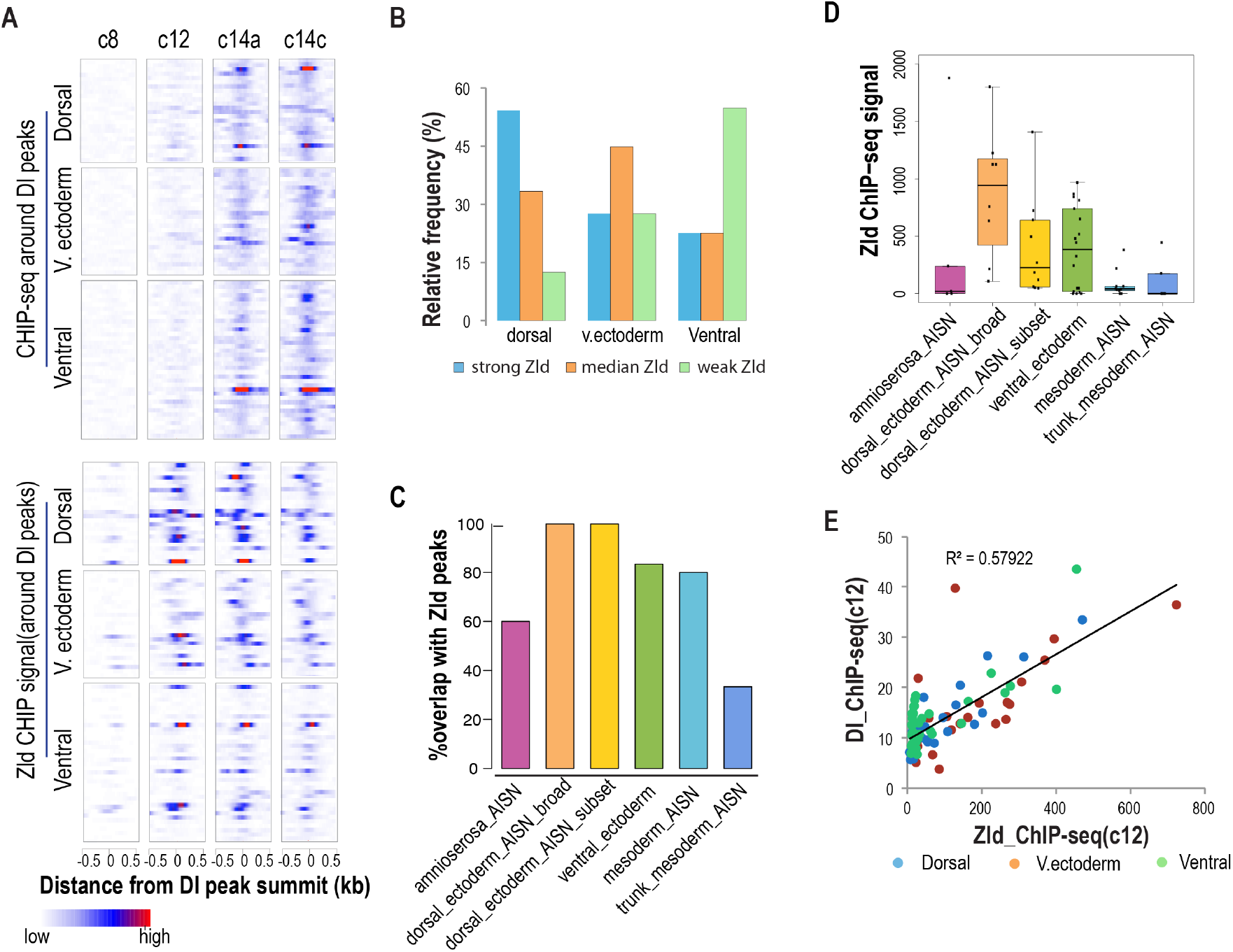
Dl and Zld binding dynamics around genes expressed in stereotypical dorsal - ventral patterns. A. Heatmaps showing the dynamics of Dl and Zld ChIP-seq signal around Dl peaks located in the intergenic and intron regions within +/− 15 kb around the dorsal – ventral genes that display primarily ventral(mesoderm), ventral ectoderm, or dorsal (including dorsal ectoderm and amnioserosa) pression pattern. These genes were selected based on published studies (Leptin, 1991; Sandmann et al., 2007; Stathopoulos et al., 2002) and the gene expression patterns annotated in the BDGP in situ image database (Hammonds et al., 2013; Tomancak et al., 2007) (http://insitu.fruitfly.org/cgi-bin/ex/insitu.pl) (Table S1). B. The plot shows the relative frequency (as percentage) of Dl peaks associated with each class of genes described in A that are either bound strongly, modestly, or weakly/not bound by Zld. For this analysis, all the Dl peaks associated with the dorsal-ventral genes were divided into three equal sized groups that displayed strong, modest, or weak/no Zld binding at cycle 12, and the relative distribution of the Dl peaks associated with each class of genes between these three groups was then calculated. C. Percentage overlap of the Vienna-tiles enhancers(Kvon et al., 2015) that drive different dorsal-ventral expression patterns with Zld peaks detected at cylce 12. Only the enhancers that were bound by Dl were used in this analysis. D. Box plots showing the relative distributions of Zld ChIP-seq signals (c12) over the Vienna-tiles enhancers belonging to different groups based on expression pattern. E. Scatter plot showing the correlation between the Dl vs Zld ChIP-seq signal around the Dl peaks associated with the three different classes of genes described in A.

To provide further evidence that Zld is likely to be more important for Dl target enhancer activity in the dorsal and lateral part of the embryo, we analyzed the association of Zld peaks with a large set of embryonic enhancers previously identified in a large-scale analysis of full-genome, non-coding sequences (Kvon et al., 2015). We focused on enhancers that drove expression with dorsal – ventral patterns, and were also bound by Dl. We found that a high percentage of dorsal and ventral ectoderm enhancers were associated with Zld peaks (Fig.3C). Among the amnioserosa enhancers, a smaller percentage were associated with Zld peaks, which may be explained by their dependence on alternative activators other than Dl for activation.

There are two sub-categories of mesoderm enhancers: mesoderm and trunk mesoderm. A relatively low percentage of the trunk mesoderm group enhancers overlapped with Zld peaks, but a large portion of the mesoderm group enhancers did. Importantly Zld signals associated with both subgroups of mesoderm are in general very low (Fig. 3D, Fig. S2), in contrast to the dorsal and ventral ectoderm enhancers. Taken together, these analyses provide further evidence that enhancers active in the dorsal, and to a lesser extent the lateral part of the embryo are much more likely to be associated with strong Zld binding than those active in the mesoderm.

Even though Dl peaks associated with mesoderm genes are less likely to be associated with strong Zld binding, some of them do. We found that these genes also displayed stronger Dl binding in early embryos than those expressed in the mesoderm but not bound by Zld, similar to Dl peaks of other two classes of genes (Fig. 3E, Fig.S3). Thus, even in the ventral part of the embryo where the Dl concentration is the highest, Zld is still required for strong Dl binding in early embryo, which presumably is important for early onset of the target gene expression.

### Zld binding and the timing of the onset of zygotic gene expression

After showing that Zld binding can affect the temporal dynamics of Dl binding, we wondered how Zld binding is related to the timing of dorsal-ventral gene expression. We focused on the genes that are expressed zygotically with no maternal expression, and ranked them according to their transcription onset time based on a previously published single embryo RNA-seq data (Lott et al., 2011) (Fig. S4A). We looked at ChIP-seq signals of Zld and Dl at the promoters of these genes (Fig. S4B), as well as at the Dl peaks in their intronic and intergenic regions (Fig. S4C). We found that early but not late expressed genes are associated with strong Zld binding at their promoters, as was shown previously for zygotic genes in general (Harrison et al., 2011). Interestingly, the later expressed genes often have strong Zld binding outside the promoter. This result shows that some zygotic genes are not expressed early, even if they are associated with strong Zld binding. For these genes, the strong binding of Zld is best explained by its requirement for boosting the Dl activity in lateral and dorsal part of the embryo where Dl concentration in the nuclei is low, as describe above. Taken all these analysis together, there are two distinct functions of Zld in coordinating the spatial and temporal enhancer activities in the early zygotic gene activation, as suggested by previous studies (Foo et al., 2014; Hannon et al., 2017; Nien et al., 2011; Xu et al., 2014).

### The relationship between Cad and Zld binding

We next analyzed the temporal dynamics of Cad binding and how it relates to Zld binding. Similar to Dl peaks, Cad peaks fall into two groups, an early group which includes peaks detected in cycle 12, and a late group which includes peaks detected only in cycle 14 (Fig. 4A). For peaks in the early group, there was, on average, stronger binding by Zld (Fig.4B), and Zld binding can be seen to precede Cad binding (Fig.4B and 4C). In addition, there is a strong correlation between Zld and Cad in early embryos (Fig. 4D), suggesting Zld may play a broad, quantitative role on Cad binding, similar to Dl. The correlation between Zld and Cad binding is diminished at cycle 14, presumably as a result of increased role of other factors in promoting Cad binding.

**Fig.4.**
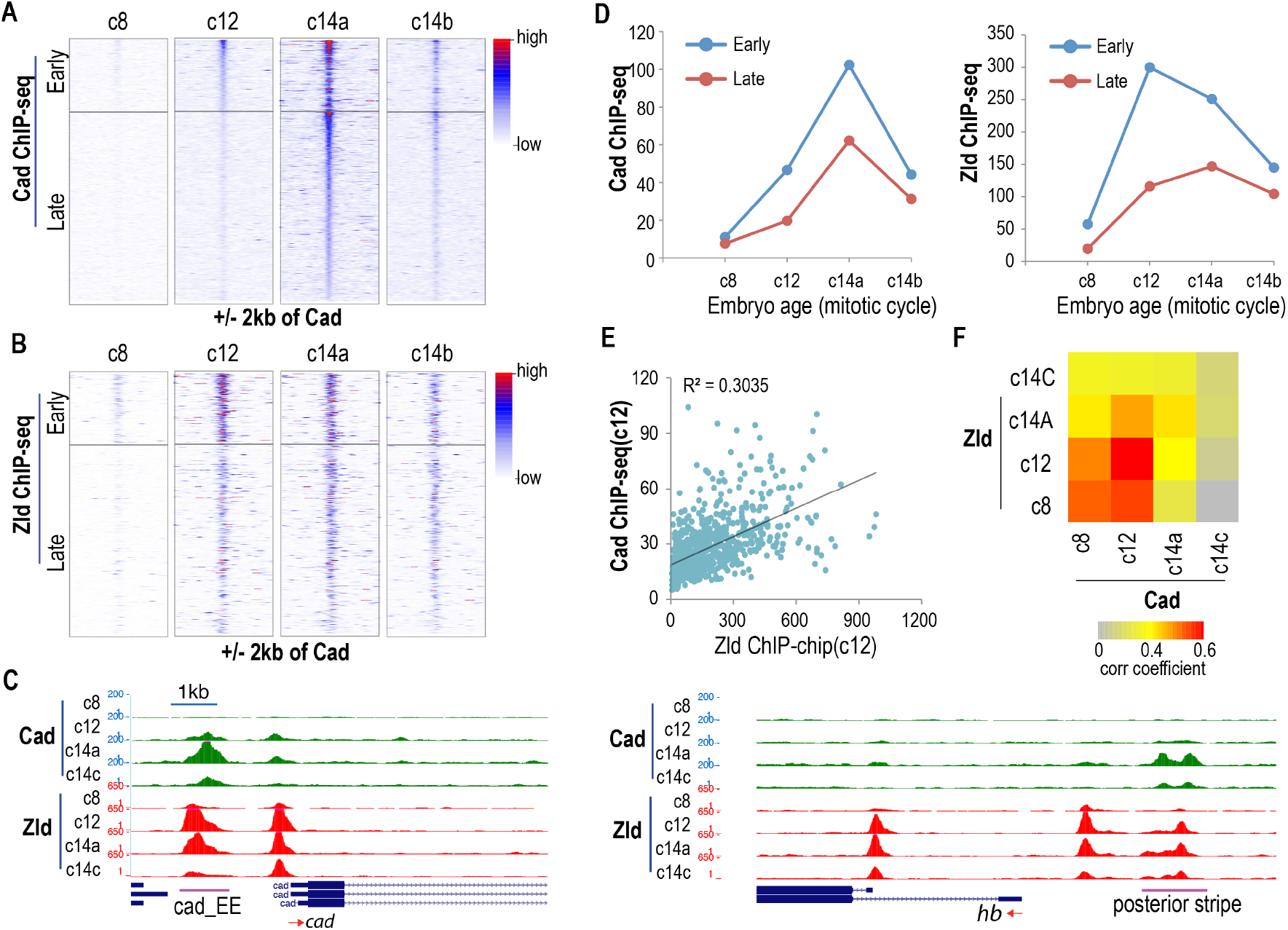
Correlation between Zld binding and the temporal dynamics of Cad binding. A. Heatmap of Cad ChIP signals around Cad peaks. The Cad peaks were divided into two groups: the early group corresponds to those that were detected in cycle 12, and the late group includes all the rest. The early binding peaks were sorted according to ChIP-seq signal strength at cycle 12, and the late binding peaks were sorted according to ChIP-seq signal at cycle 14a. B. The Zld ChIP signals around the two groups of Cad peaks. The peaks were sorted as in A. C. Genome browser snapshots of two Cad peak regions, including the putative *cad* embryonic enhancer and the *hb* posterior stripe enhancer (Margolis et al., 1995), showing the dynamics of Cad and Zld ChIP-seq signal. D. Overall trend for the Cad and Zld signal for the two groups of Cad peaks. E. Correlation between Zld and Cad signals around Cad peaks of both groups at cycle 12. F. Heatmap showing correlation coefficient between Cad and Zld ChIP signals at different stages.

The Cad activated genes have been less as well studied as the Dl activated genes. This precludes similar detailed analyses of the spatial effect of Zld on Cad dependent genes as described above for Dl dependent genes. Nevertheless, based on a Zld binding site mutation analysis using transgenic reporter assay we found that Zld also functions to boost Cad target enhancer activity in cells with low Cad concentrations (not shown), suggesting it also important for both spatial and temporal effect on Cad target enhancer activity and target gene expression.

### Zld binding in the early embryo leads to deposition of histone acetylation marks and local nucleosome depletion

In a previous study, we showed that genome wide Zld binding is highly correlated with deposition of histone acetylation marks in early embryos, and that these histone marks were detected as early as cycle 8, when the enhancers are not thought to be functional (Li et al., 2014). This suggested that Zld might play a direct role in recruiting histone modifying enzymes to those loci. However, we could not exclude the possibility that binding of other factors might also be important. Now we have shown Zld binding occurs earlier than Dl and Cad in regions cobound by either of these factors, we wondered whether the histone acetylation marks are detectable upon Zld binding. Significantly, as shown in Fig. 5A-B, we found that the early Dl and Cad binding peaks, which are generally associated with strong Zld binding, were already associated with relatively strong histone acetylation signals at cycle 8, when Zld but not Cad or Dl was bound. Together with the near perfect correlation between early Zld binding and the presence of Zld binding sites (Harrison et al., 2011), this result provides strong evidence that Zld, and not the maternal factors it helps to recruit, is likely to be directly responsible for the deposition of these histone marks in the Dl and Cad peak regions at cycle 8.

**Fig.5.**
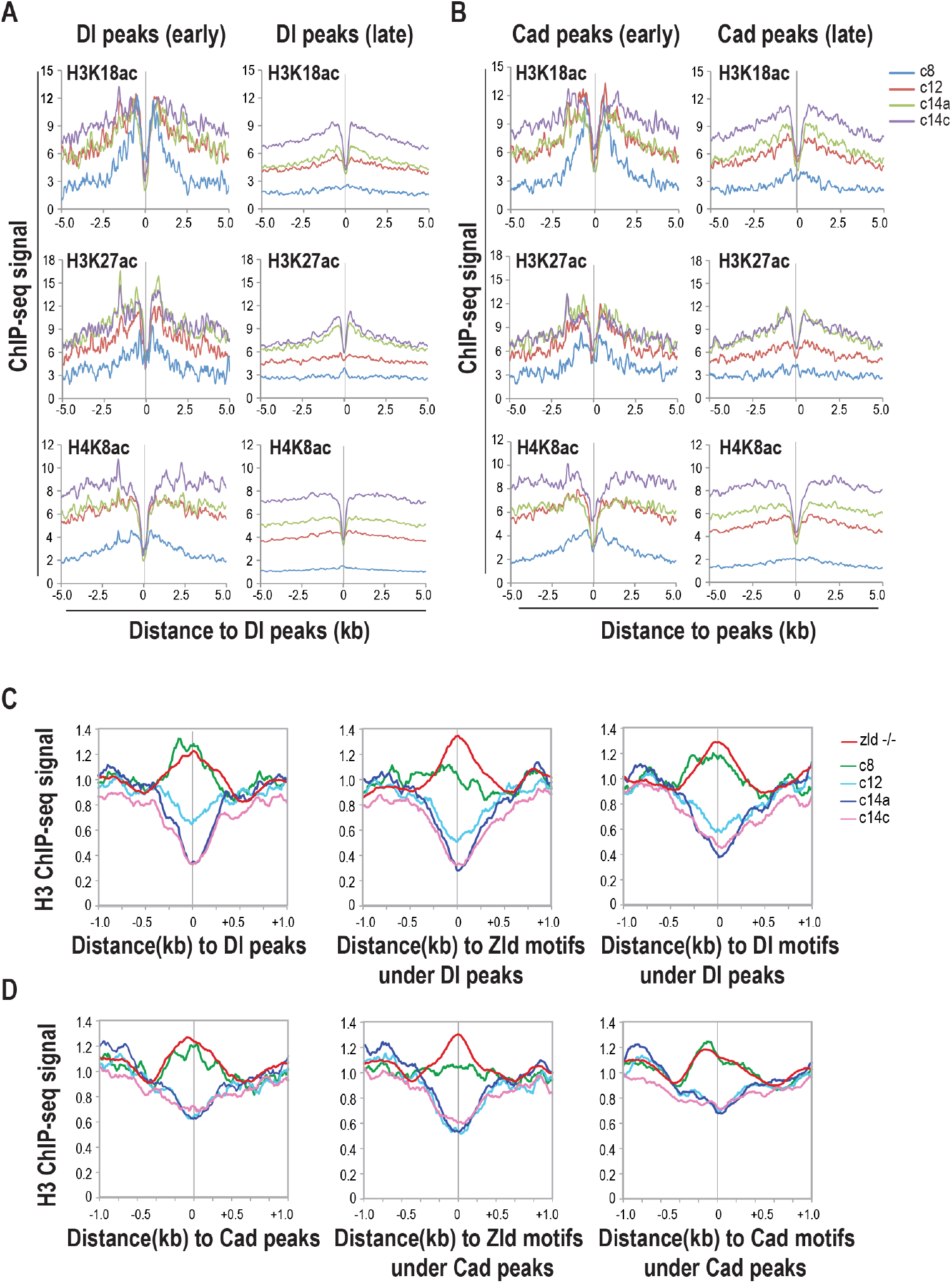
Zld binding leads to deposition of histone acetylation marks and localized histone depletion prior to Cad and Dl binding. A. Composite ChIP-seq signal profiles for histone acetylation marks based on previously reported data (Li et al., 2014) around Dl peaks at different stages are shown. The Dl peaks are divided into early and late binding groups as described in Fig. 2A. B. Similar to A except histone acetylation profiles around early and late Cad peaks, grouped as described in Fig.4A, are shown. C. Histone H3 ChIP-seq signal profiles around early Dl peak center, or the Dl and Zld motifs located within 300 bp of those peaks in cycle 8, cycle 12, cycle 14a and cycle 14c embryos are shown. The ChIP-seq signal profiles from *zld*- embryos were included for comparison. D. Similar to C, except that the profiles around early Cad peaks, and the Cad and Zld motifs in those peak regions are shown.

We also wondered whether Zld binding by itself can cause nucleosome depletion. Based on histone H3 ChIP-seq analysis in our previous study (Li et al., 2014), we showed that histone depletion at zygotic enhancers was not apparent at cycle 8. However, we reasoned that there might be more localized histone depletion at Zld binding sites that escaped previous analysis. Therefore, we carried out more detailed analysis to include not only the histone ChIP-seq signals centered at Dl and Cad peaks, but also around their motifs and the Zld motifs located in those peak regions. We limited our analysis to the Dl and Cad peaks that were detected at cycle 12 since these loci were bound by Zld early and more strongly. We compared these signals to those detected in *zld*- embryos lacking Zld. As we showed previously, histone depletion at zygotic enhancers is broadly affected in the *zld*- embryos, and thus the histone H3 ChIP signal in these embryos can be considered to represent the inactive state.

From the above analysis, we found that the histone H3 ChIP signal centered around Dl (Fig.5C) and Cad (Fig.5D) peaks at cycle 8 is comparable to the signal from *zld*- mutant embryos, suggesting an overall lack of histone depletion at this early stage. Interestingly, the signal around the Zld motifs is much lower at cycle 8 compared to the mutant embryo, even though the signal around the Dl and Cad motifs did not show a significant change. This result suggests that Zld binding at cycle 8 caused local histone depletion over the Zld motifs in the Dl and Cad peak regions before Dl and Cad binding was detected.

### Zld binding in coding regions led to histone acetylation and increased chromatin accessibility

The analyses above suggest that Zld has a direct role in histone modification and depletion. To provide further evidence, we carried out analyses on a group of Zld binding peaks located in the coding regions (CDS) of the genome, which most likely represent non-functional Zld binding, independent of other factors.

Previously, we showed that a surprisingly large fraction of Zld motifs were bound by Zld in cycle 8 embryos at all genomic regions, including the CDS (Harrison et al., 2011). Zld binding in the CDS was as strong as in other genomic regions in early embryos, but decreased rapidly at later time points, an effect reflected in our current ChIP-seq data, as shown in Fig.S5A. This represents the first evidence that Zld binding in CDS is likely to be non-functional. Further, by focusing on the top 1000 Zld peaks ranked based on binding at cycle 12, we found that Zld peaks in the CDS, compared to Zld peaks located in intergenic and intron regions, are much less likely to overlap with other transcription factors present in the early transcription network that have previously been studied by ChIP-chip or ChIP-seq (Fig.S5B). For the Zld peaks in the CDS that do overlap with binding of other factors, the binding levels of those factors are also much lower than Zld peaks in other types of sequences (e.g. intergenic and intron regions, Fig. S5C). All these observations strongly indicate that the Zld peaks in the CDS represent non-functional binding and are co-bound by few other factors.

Interestingly, in spite of likely being non-functional, we found that Zld peaks in the CDS are nevertheless associated with histone acetylation marks (Fig.6A). In addition, based on comparison to data from *zld*- embryos, there is also significantly lower histone H3 ChIP-seq signal around the CDS Zld peaks in early embryos (Fig.6B), suggesting that Zld binding in the CDS can cause local histone depletion.

**Fig.6.**
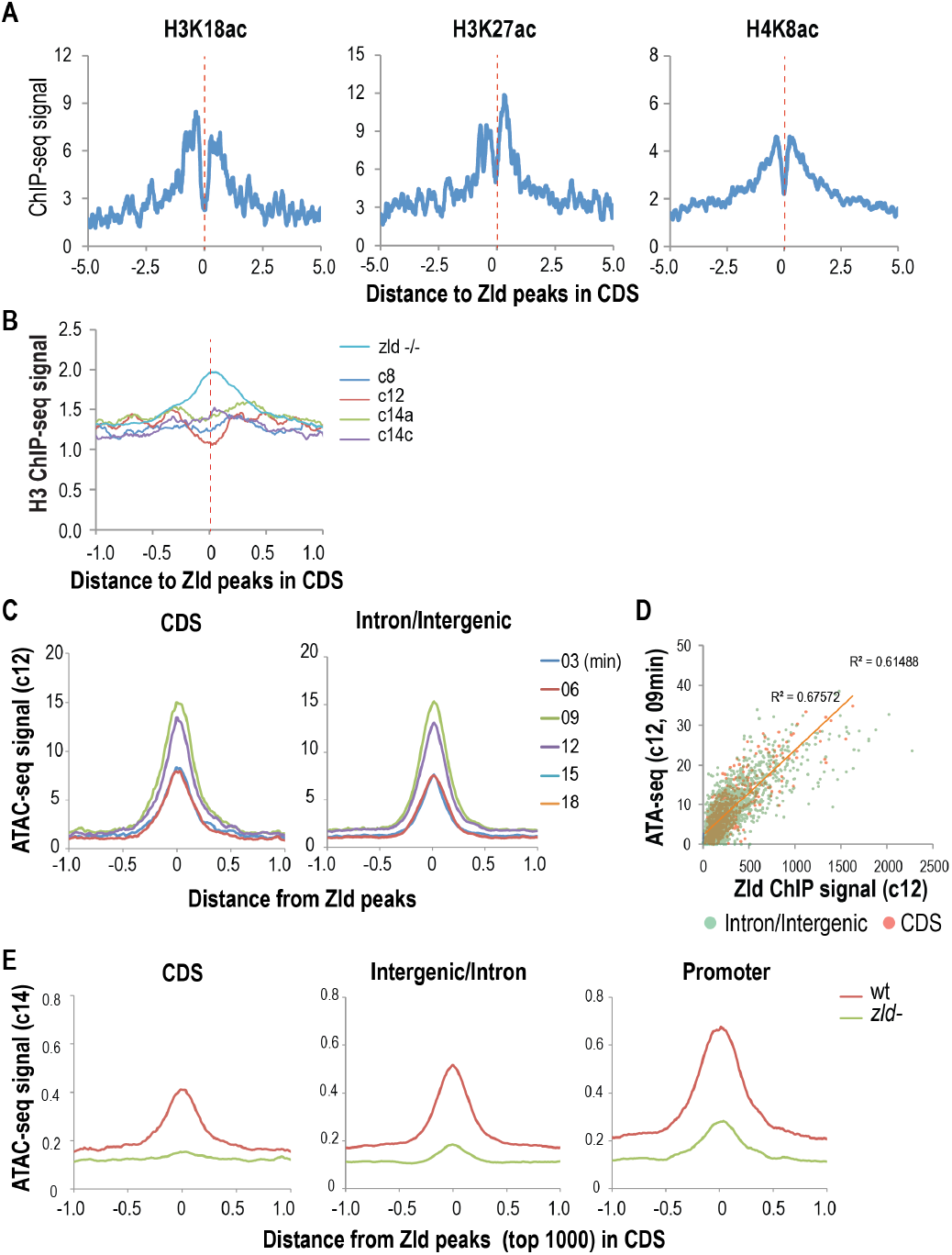
Zld binding in coding sequences leads to the deposition of histone acetylation marks, histone depletion, and increased chromatin accessibility. A. Dynamics of the ChIP-seq signal profile of histone acetylation marks around of Zld peaks (top 1000 based on signal at cycle 12) located in the CDS. B. Dynamics of the histone H3 ChIP-seq signal profile. The ChIP-seq signal profile in *zld-* mutant embryos (Li et al., 2014) is also shown. C. The dynamics of ATAC-seq signals in mitotic cycle 12 around the Zld peaks located in the CDS, or intergenic and intron sequences are shown based on data published previously(Blythe and Wieschaus, 2016). D. Correlation between accessibility and ChIP signal of Zld peaks located in the CDS, and those located in intergenic and intron regions. E. The chromatin accessibility around the Zld peaks in the CDS, the intergenic and intron regions, as well as the promoter decreased dramatically in zld-mutant embryos based on data from a previous study(Hannon et al., 2017).

To further investigate the effect of Zld binding in the CDS on chromatin, we analyzed the chromatin accessibility around the CDS peaks using single embryo ATAC-seq data previously published for cycle 11 – cycle 13 embryos (Blythe and Wieschaus, 2016). We found that Zld peaks in the CDS are associated with strong ATAC-seq signals, at levels comparable to what is observed around Zld peaks located in the intergenic and intron regions as shown in Fig. 6C in cycle 12 embryos. Importantly, Zld binding in the CDS, as well as intergenic and intronic regions, showed a similar quantitative relationship to the ATAC-seq signal (Fig.6D).

To show that Zld binding is directly responsible for the chromatin accessibility around the Zld peaks in CDS, we compared the ATAC-seq signals in wild type and *zld-* embryos based on published data collected in early c14 embryos (Hannon et al., 2017), and found that the ATAC-seq signal around Zld peaks in the CDS is completely abolished in the *zld-* mutant embryos (Fig.6E). Taken together, Zld binding to the CDS led to deposition of histone marks, histone depletion and increased chromatin accessibility as in other sequences. Since the CDS peaks were not bound or only bound weakly by other factors, these observations provide further evidence that Zld binding can directly cause these changes in chromatin structure.

### Zld binding has stronger impact on chromatin accessibility than Cad and Dl

Active enhancers are known to be located in open chromatin regions, and thus transcription factor binding is expected to correlate with chromatin accessibility. Interestingly, we found that while Dl, Cad, as well as Zld ChIP-seq signals correlated with ATAC-seq signals, the correlation for Zld is much stronger, as shown in Fig. S6A,B,E,F in cycle 12 embryos, suggesting Zld binding can have much stronger effect on chromatin accessibility. To investigate this further, we divided the early group of peaks for Cad and Dl into three subgroups based on Zld ChIP-seq signal. We found that the levels of Cad and Dl binding between the subgroups are on average similar (Fig. S6C, and S6G), which can be attributed to the presence of a larger number of/stronger Cad and Dl motifs around the peaks in the weak-Zld subgroups(not shown). We found that the peaks in the strong-Zld subgroup for each factor are associated with much higher ATAC-seq signal (Fig. S6D, and S6H), confirming that Zld binding can have much stronger effect on chromatin accessibility than Cad and Dl.

### Zld motifs are associated with positioned nucleosomes

Zld bound regions and early zygotic enhancers have been shown to be associated with high nucleosome occupancy (Sun et al., 2015), which has been proposed to be important for enhancer function. Interestingly, we observed that in *zld-* embryos, the histone H3 ChIP-seq signal is much higher around Zld peaks in the CDS (Fig.6B), suggesting that non-enhancer Zld peaks are also associated with higher nucleosome occupancy. To provide further evidence for this, we analyzed the nucleosome occupancy profile based on MNase-seq data generated in S2 cells (Chereji et al., 2016), in which Zld is not expressed. As shown in Fig. 7A, the nucleosome signal is higher around Zld peaks in the CDS regions even if the flanking sequences are already associated with much higher MNase-seq signal than the Zld peaks in intergenic and intron regions. Furthermore, when we analyzed the distribution of the MNase-seq read fragment midpoints around Zld motifs in the top 1000 Zld peaks (ranked based on ChIP-seq signal in cycle 12), we saw a strong enrichment in the +/− 70 bp region around the motifs, for both CDS and the intergenic and intron peaks (Fig. 7B),. Surprisingly, when we analyzed the unbound Zld motifs located in the intergenic and intron, or matching CDS regions, we observed positioned nucleosomes over Zld motifs as well, even though the nucleosome levels over these unbound motifs are only modestly higher than flanking sequences. This suggests that strong *a priori* nucleosome positioning is a general feature of Zld motifs.

**Fig.7.**
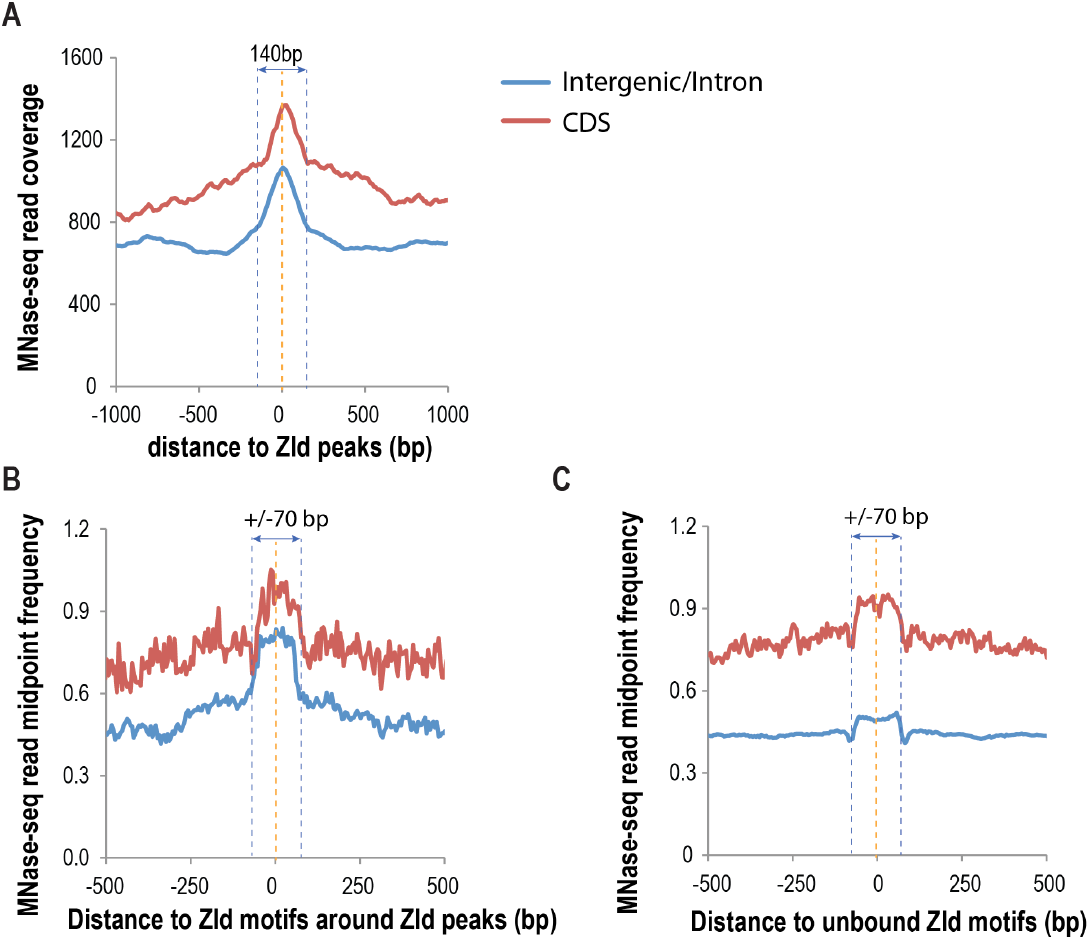
High nucleosome occupancy and nucleosome positioning around Zld peaks and motifs. A. MNase-seq signal profile around Zld peaks (top 1000 ranked based on signal in cycle 12) that are located in the CDS, or intergenic and intron regions. The MNase-seq data was obtained from S2 cells (Chereji et al., 2016). B. The distribution of MNase-seq read midpoint around Zld motifs located in the Zld bound regions in the CDS or intergenic and intron sequences. C. The distribution of MNase-seq read midpoint around Zld motifs located in the CDS or intergenic and intron sequences that were not bound Zld.

## Discussion

### Effect of Zld binding on chromatin structure

The eukaryotic genome is packed into chromatin through formation of the nucleosomes, which can sterically occlude the DNA recognition sequences from recognition and binding by transcription factors (Kornberg and Lorch, 1999) (Khorasanizadeh, 2004; Liu et al., 2006). Even though active enhancers are located within accessible chromatin (John et al., 2008; Li et al., 2011; Nègre et al., 2011; Sabo et al., 2004; Song et al., 2011; Thurman et al., 2012), they have been shown, both experimentally and computationally, to be associated with high nucleosome occupancy (Barozzi et al., 2014a; Sun et al., 2015; Tillo et al., 2010) when they are not active. An important question is how transcription factors gain access to enhancers in closed chromatin to effect transactivation. An important, emerging mechanism is pioneer factors(Zaret and Carroll, 2011).

A distinguishing characteristic of pioneer factors is their ability to bind to enhancers in closed chromatin, subsequently making enhancers accessible and promoting binding by other factors(Zaret and Carroll, 2011). Previous studies have shown that Zld is important for histone depletion and chromatin accessibility at its target enhancer regions(Foo et al., 2014; Li et al., 2014; Schulz et al., 2015; Sun et al., 2015). In addition, Zld binding is correlated with deposition of histone acetylation marks in early embryos(Li et al., 2014). However, it was not clear whether Zld binding itself can change chromatin state.

In this study, we found that the binding of Zld in the peak regions of two major maternal factors, Cad and Dl, led to deposition of histone acetylation marks, and local histone depletion around Zld motifs at cycle 8 before Dl and Cad binding was detected. In addition, we showed that even though the Zld peaks in the CDSs were not co-bound or were bound at very low levels by other transcription factors in contrast to the intergenic and intronic peaks, they were also associated with histone acetylation and depletion, as well as increased local chromatin accessibility as measured by ATAC-seq; and the Zld ChIP-seq signal and ATAC-seq signal at Zld peaks in the CDS are also strongly correlated, similar to Zld peaks in intergenic and intron regions at cycle 12. Together, these results strongly suggest that Zld has a direct role in increasing histone acetylation and local chromatin accessibility around its binding sites, presumably by recruiting histone acetyltransferases and nucleosome remodeling complexes to the enhancers.

### Functional roles of the Zld on the temporal dynamics of transcription factor binding and enhancer activities along morphogen gradients

Zld binding broadly overlaps with other factors in the early embryo transcription network (Harrison et al., 2011). In this study, we showed that the binding of Dl and Cad correlated quantitatively with Zld binding, and as a result Zld can have a broad, quantitative effect on the binding of these factors, and by extension the activities of the enhancers targeted by these factors.

The effects of Zld on enhancer activities can be attributed to two distinct functions. One of them is to boost activity of patterning transcription factors in cells with low factor concentrations along the morphogen gradients as previously proposed(Foo et al., 2014) and demonstrated for Bcd enhancers (Hannon et al., 2017; Xu et al., 2014). In this study, we extended the previous studies to the Dl target enhancers. We showed that enhancers that function in the dorsal and lateral part of the embryo are much more likely to be associated with strong Zld binding than enhancers that are active along the ventral/mesoderm part of the embryo, where Dl concentration is high. At the same time, we showed that Zld can promote early binding of Dl and Cad, which presumably is important for early onset of some of the target genes during zygotic genome activation. In support of this, a previous study showed that the expression of some genes appeared to be delayed in *zld*- mutant embryos (Nien et al., 2011).

Based on our study, the effects of Zld on the temporal and spatial enhancer activity are distinguishable. We showed that for Dl activated genes, even though the dorsal and lateral enhancers are more likely to be associated with strong Zld binding, they do not necessarily display early onset in transcription, which requires binding of Zld at the promoter. Thus, in such cases the association with strong Zld binding may be attributed to requirement for Zld to promote Dl binding in the presence of low Dl concentration. On the other hand, for the ventral/mesoderm enhancers, even though most of them are not associated with strong Zld binding, some of them do. For these enhancers, the strong association with Zld may be best explained for its requirement for early transcription onset for these genes. Taken together, Zld serves as integral part of the early transcription factor network important for determining the precise spatiotemporal patterns of zygotic gene expression.

Pioneer factors in other systems have been shown to serve diverse functions. The role of Zld in promoting early transcription factor binding is reminiscent to the pioneer factor PHA-4 in *C. elegans*. The genes activated by this factor during the pharyngeal development are expressed in an ordered fashion that is determined by the strength of their PHA-4 binding sites (Gaudet and Mango, 2002). Other pioneer factors are often found to bind to enhancers prior to target gene expression during cell fate specification during development (Iwafuchi-Doi and Zaret, 2016). For example, the first pioneer factors identified, FoxA and GATA, bind to a liver specific gene, Abl1, in liver progenitor cells before liver tissue induction (Gualdi et al., 1996). Such early binding or pre-marking of enhancers may be required to prevent the formation of repressive chromatin structure and maintain functional competence (Xu et al., 2009). Moreover, pioneer factors have been shown to prime enhancers prior to hormonal induction (Zaret and Carroll, 2011). Our data add to a growing body of data that suggest that pioneer and pioneer like transcription factors play diverse roles in regulating enhancer function.

### Nucleosome positioning over Zld motifs

Previous studies have indicated that enhancers and transcription factor bound sequences are usually associated with high nucleosome occupancy (Ballaré et al., 2013; Barozzi et al., 2014a; Sun et al., 2015; Tillo et al., 2010). In particular, previous studies on Zld (Sun et al., 2015) and the pioneer factor Pu.1(Barozzi et al., 2014b), which is associated with hematopoietic lineage specification, have shown that they often bind to sequences that are associated with high nucleosome occupancy, which are determined by the sequences flanking the motifs of these factors. It has been suggested that high nucleosome occupancy is tightly related to the binding of these two factors and enhancer function. However, in this study we showed that higher nucleosome occupancy is not limited to Zld binding sites in the enhancer-rich intergenic and intron regions. Instead, the Zld binding sites in the CDS, which are likely to be non-functional, and even unbound Zld motifs, are also associated with higher nucleosome levels. A careful examination of data from the previous study on Pu.1 shows that the unbound Pu.1 motifs were associated higher nucleosome occupancy too, albeit at more modest levels, similar to what we observed with Zld.

Thus, higher nucleosome levels associated with transcription factor motifs, particularly for pioneer factors such as Zld and Pu.1, may be a general phenomenon, instead of uniquely related to enhancer or transcription factor binding sequences. Importantly, not only the Zld and Pu.1 motifs are associated with higher nucleosome occupany, for both factors, there is much higher MNase-seq fragment midpoint density within a ~ +/− 70 bp region around their motifs, suggesting strong nucleosome positioning surrounding the motifs.

It is not clear why enhancer sequences, and in particular the sequences associated with Zld and Pu.1 motifs, are associated with high nucleosome occupancy. Based on our findings, we favor the a model (Sun et al., 2015) in which nucleosome occupancy is important for preventing non-functional binding of transcription factors to chromatin. At enhancers, where there is strong enrichment of transcription factor binding sites, high nucleosome occupancy may be important to prevent non-functional binding in cells where an enhancer is no longer active. In the case of Zld, strong, non-functional binding does occur in the CDS, and presumably in other non-enhancer sequences in early embryos, where the chromatin is highly accessible. However, as the embryo enters cycle 14, Zld binding in the CDS decreased dramatically, at much faster pace than observed for the intergenic and intron peaks. This may be explained at least in part by the much higher nucleosome occupancy in the CDS, consistent with a role of high nucleosome occupancy in inhibiting non-functional binding.

An alternative hypothesis is that high nucleosome occupancy at enhancers may be necessary for their function. However, we believe this is unlikely as it is well known that juxtaposition of nucleosome destabilizing sequences next to transcription factor binding sites can actually promote enhancer activity, as exemplified by the poly(dA-dT) tract associated with yeast promoters (Iyer and Struhl, 1995; Raveh-Sadka et al., 2012). The study on Pu.1 also showed that among the Pu.1 bound regions, sequences with low nucleosome occupancy were actually more highly bound by Pu.1 in vivo, and high nucleosome density led to low Pu.1 binding at least in vitro(Barozzi et al., 2014b).

How transcription factors overcome high nucleosome occupancy and gain access to enhancers is not well understood. Pioneer factor activity provides one possible mechanism even though how such factors achieve selective binding to the enhancers remains unknown. Enhancers usually contain not only the binding sites of different factors necessary for combinatory control of transcription, but also multiple copies of binding sites for the same factors. Such high density of transcription factor binding sites may allow transcription factors to outcompete nucleosomes through cooperative competition (Miller and Widom, 2003; Mirny, 2010). Thus, transcriptional regulatory specificity appears to be achieved through a combination of the association of transcription factor motifs with high nucleosome occupancy and the countering effect of collective binding of multiple factors at enhancers.

Based on previous analyses of the Pu.1 and Zld binding sequences, nucleosome occupancy is determined by sequences flanking the motifs of these factors, instead of the motif sequences themselves (Barozzi et al., 2014a; Sun et al., 2015). Since sequence features determining nucleosome affinity and positioning may be more extensive and diffused, the observed association of motifs with nucleosomes may be best explained by the elimination of transcription factor motifs that are not covered by nucleosomes, instead of change of nucleosome positioning that allow overlap of nucleosome and factor motifs during evolution.

## Materials and Methods

### Embryo collection, fixation with formaldehyde, chromatin preparation

Flies were maintained in population cages, embryos collected, crosslinked in vivo with formaldehyde, and manually sorted to obtain embryos that distributed around cycle 8, 12, 14a, and 14c. The chromatin was prepared through CsCl ultracentrifugation following the same procedure previously as described in (Li et al., 2014).

### ChIP and sequencing

As previously described, the chromatin was fragmented to sizes ranging from 100 to 300 bp using a Bioruptor (Diagenode, Inc., Seraing, Belgium), and each sample was mixed with a roughly equivalent amount of chromatin isolated from stage 4/5 (mitotic cycle 13-14) *D. pseudobscura* embryos. The chromatin immunoprecipitation reactions were carried out as described previously (Harrison et al., 2011). For each reaction, we used 2 μg of the mixed chromatin, and either 1 μg of anti-Bcd(1), 1 μg of anti-Dl(3), 1 μg of anti-Cad(2), or 1 μg of anti-Zld antibody as described in (Harrison et al., 2011; Li et al., 2008). The sequencing libraries were prepared from the ChIP and Input DNA samples using the Illumina (San Diego, CA) TruSeq DNA Sample Preparation kit following the manufacturer’s instructions, and DNA was subjected to ultra-high throughput sequencing on a Illumina HiSeq 2000 DNA sequencer.

### Mapping sequencing reads to the genome, and peak calling, and signal normalization

With read length of 100, the sequenced reads were mapped jointly to the April 2006 assembly of the *D. melanogastergenome* [Flybase Release 5] and the November 2004 assembly of the *D. pseudoobscuragenome* [Flybase Release 1.0] using Bowtie(Langmead, 2010) with the command-line options ‘-q -3 50 -n 2 -l 50 -a -m 1 --best –strata’ to ensure accurate mapping of reads to *D.melanogaster* and *D. pseudoobscura* genomes. We called peaks for each experiment using MACS(Zhang et al., 2008) v1.4.2 with the options ‘-c -n -g dm --nomodel --shiftsize=90’, and used Input as controls. To generate the ChIP-seq profiles, each read was extended to 180 bp based on its orientation, and the resulting profiles (in wig format were normalized based on the number of aligned reads for *D.melanogaster* and *D.pseudoobscura* as previously described (Li et al., 2014).

### Dynamics of ChIP signals

To analyze the dynamics of ChIP signals at different stages, we first generated a union peak list for each factor. Starting with peaks called by MACS as described above for each sample, we generated a consolidated list of peaks for each factor for all stages by joining each group of peaks that are within 200 bp (from summit to summit) into a single peak. We calculated the ChIP signal for each union peak at different stage by summing the ChIP signal around a 500 bp window centered around each peak position based on the normalized ChIP profile generated as described above. Similarly, to analyze the relationship between Dl and Cad binding to Zld binding, we also calculated the Zld ChIP signal around Dl and Cad peaks in a 500 bp window centered around each peak center.

### Analysis of Dl peaks around different classes of dorsal ventral genes

We generated a list of genes that are expressed predominantly along the ventral (mesoderm), lateral (v. ectoderm), or dorsal (including dorsal ectoderm and amniosoresa) domains of the embryo primarily based on the BDGP in situ image database (Hammonds et al., 2013; Tomancak et al., 2007) (http://insitu.fruitfly.org/cgi-bin/ex/insitu.pl). We first select all genes with dorsal-ventral expression pattern annotations at stage 4-6, and then visually selected genes that are primarily expressed in these three domains based on images available on the BDGP gene expression pattern website. We excluded genes that have both ventral and dorsal patterns, and genes that also have significant anterior-posterior patterns (stripes). In addition, we also added to the list genes that have previously shown to display distinct dorsal-ventral expression patterns (Leptin, 1991; Sandmann et al., 2007; Stathopoulos et al., 2002), but were not included in the BDGP database. To analyze the Dl bound enhancers around the three classes of selected genes, we identified the Dl peaks that are within +/-15 kb around each gene, and only used peaks that are in intergenic and intron regions in our analysis.

### Analysis of Dl and Zld binding to Vienna-Tiles enhancers

The Vienna-Tiles enhancers (Kvon et al., 2015) that showed expression in the “amnioserosa”, “dorsal”, “v. ectoderm”, and “mesoderm” were selected. We limited our analysis to enhancers with three or less annotated patterns. In cases with multiple terms, only the ones with “procephalic”, “endoderm”, and “hindgut” besides the dorsal-ventral pattern terms, were kept. We excluded enhancers with both dorsal and ventral patterns. In addition, enhancers annotated for both “amnioserosa” and “dorsal ectoderm” were grouped into the dorsal ectoderm groups. In addition, we only reported the enhancers that are bound by Dl in our analysis to exclude enhancer that may mainly depend on other factors for activation.

### Identification of transcription factor binding motifs

The transcription factor binding motifs in a +/-150 bp region around each transcription factor ChIP-seq peak were predicted with PATSER (version 3e) (Hertz and Stormo, 1999) with parameters set as follows: -c -s -A a:t 3 c:g 2 -lp -6. The Position Weight Matrices used for Bcd(Li et al., 2008) and Dl(MacArthur et al., 2009) and Zld (Enuameh et al., 2013)have been described previously. In the final analyses, the motifs were further filtered using a motif weight score cutoff of 3.0.

### MNase-seq profiles

We used the MNase-seq data obtained from S2 cells (Chereji et al., 2016) digested at “high” MNase concentration. Several other MNase-seq data sets in fly embryo and S2 cells have been reported, but we found this data set displayed more even genome-wide read coverage. Nevertheless, we obtained similar results as reported here when other data sets were used. In all our analyses, the MNase-seq reads were filtered based on read length and the reads of 120 – 160 bp in length were kept and used to calculate the sequence coverage profile or the MNase-seq read midpoint distribution.

### Overlap between Zld peaks in coding sequences with other transcription factors

The transcription factor binding peaks identified from previous ChIP-chip (Li et al., 2008; MacArthur et al., 2009) and ChIP-seq (Bradley et al., 2010; He et al., 2011) studies were used in the analysis. The cutoff of peak-peak distance of 500 bp was used. To show how strongly the factors were bound to the Zld binding site, the sum of the enrichment score for overlapping transcription factors for each Zld peak was calculated by adding the log2 enrichment scores for all overlapping factors in the ChIP-chip data set. Since the 21 factor ChIP-chip data set was produced under the similar experiment conditions and using the same data analysis pipeline, the sum of the enrichment scores provide a good measure of overall binding by these factors. We have not carried out similar analysis using the ChIP-seq data set since the ChIP-chip data set already include all factors in the ChIP-seq data set.

### Data availability

All raw data are available at the GEO database under the accession number.

---

**Fig. S1.**
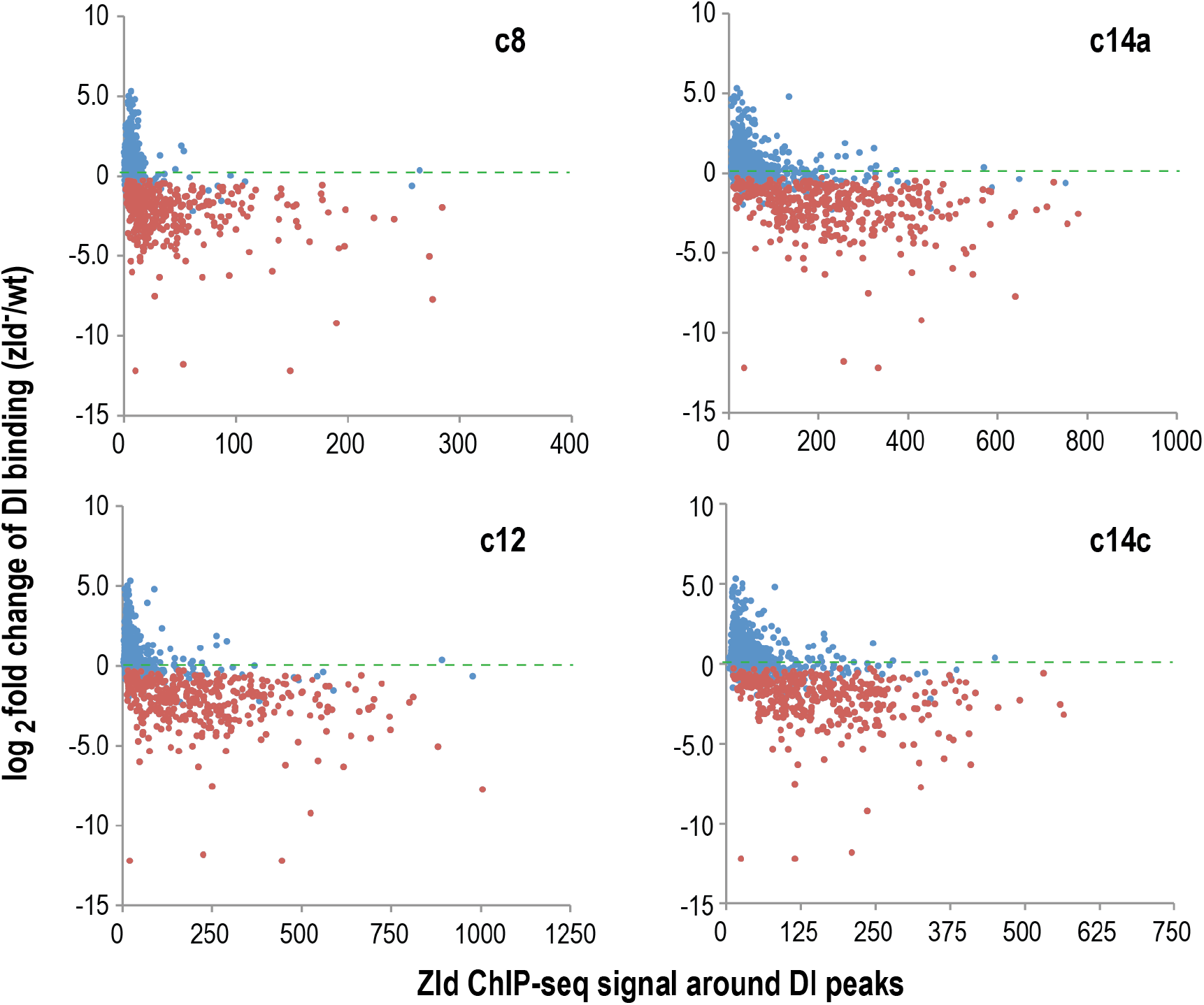
Relationship between dependence of Dl binding on Zld and the levels of Zld binding at the Dl binding sites. Similar to Fig.2G, the plots show the relationship between the magnitudes of effects of *zld* mutation on Dl binding based on data from (Sun et al., 2015) and the levels of Zld binding around the Dl binding sites detected by ChIP-seq in cycle 8, 12, c14a and 14Cembryos.

**Fig. S2.**
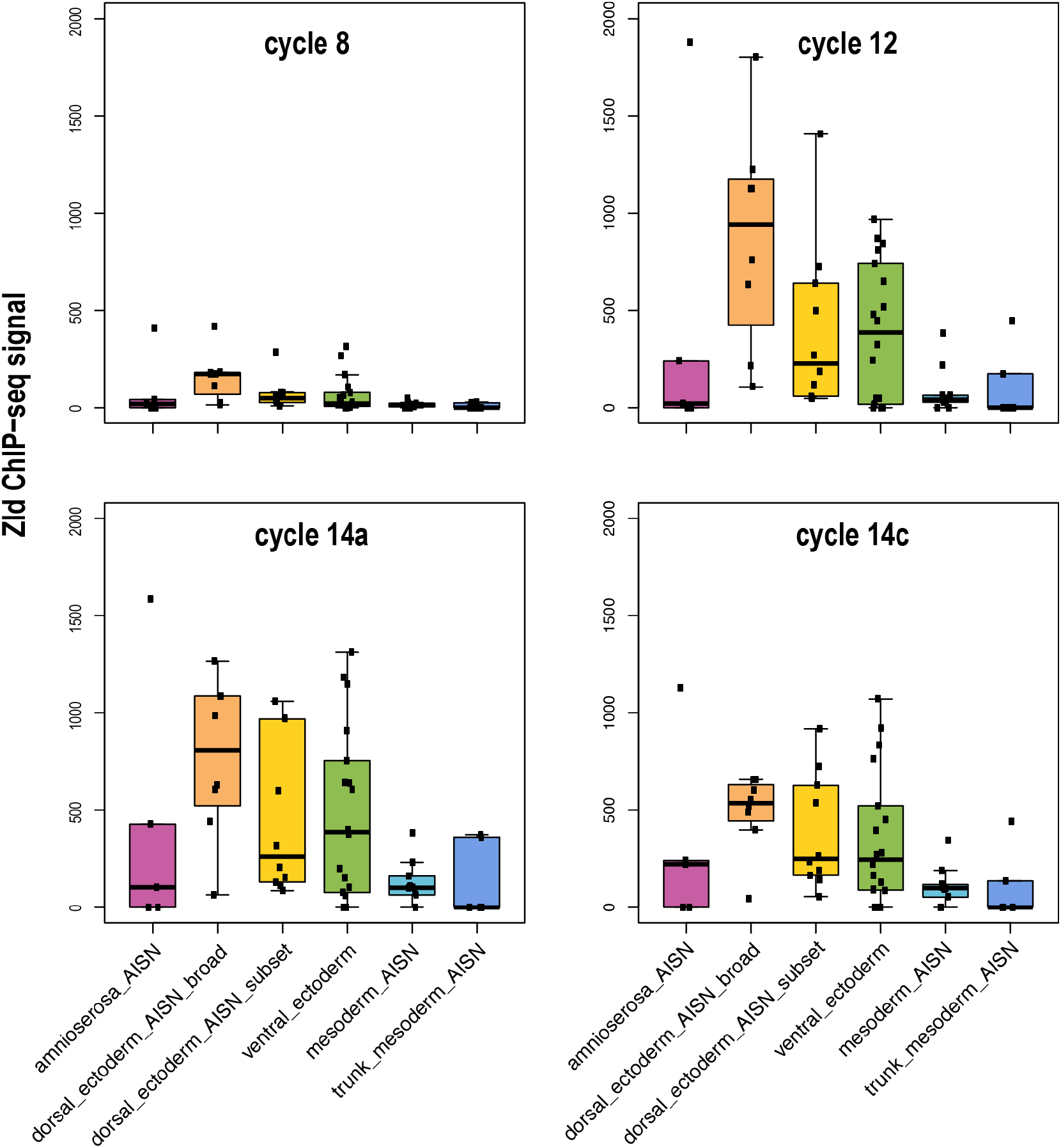
The enhancers (Kvon et al., 2015) that drive different expression patterns along the dorsal-ventral axis were bound at different levels by Zld. Similar to Fig. 2E, except that the Zld binding at cycle 8, 12, c14a, as well as cycle 14c is shown.

**Fig. S3.**
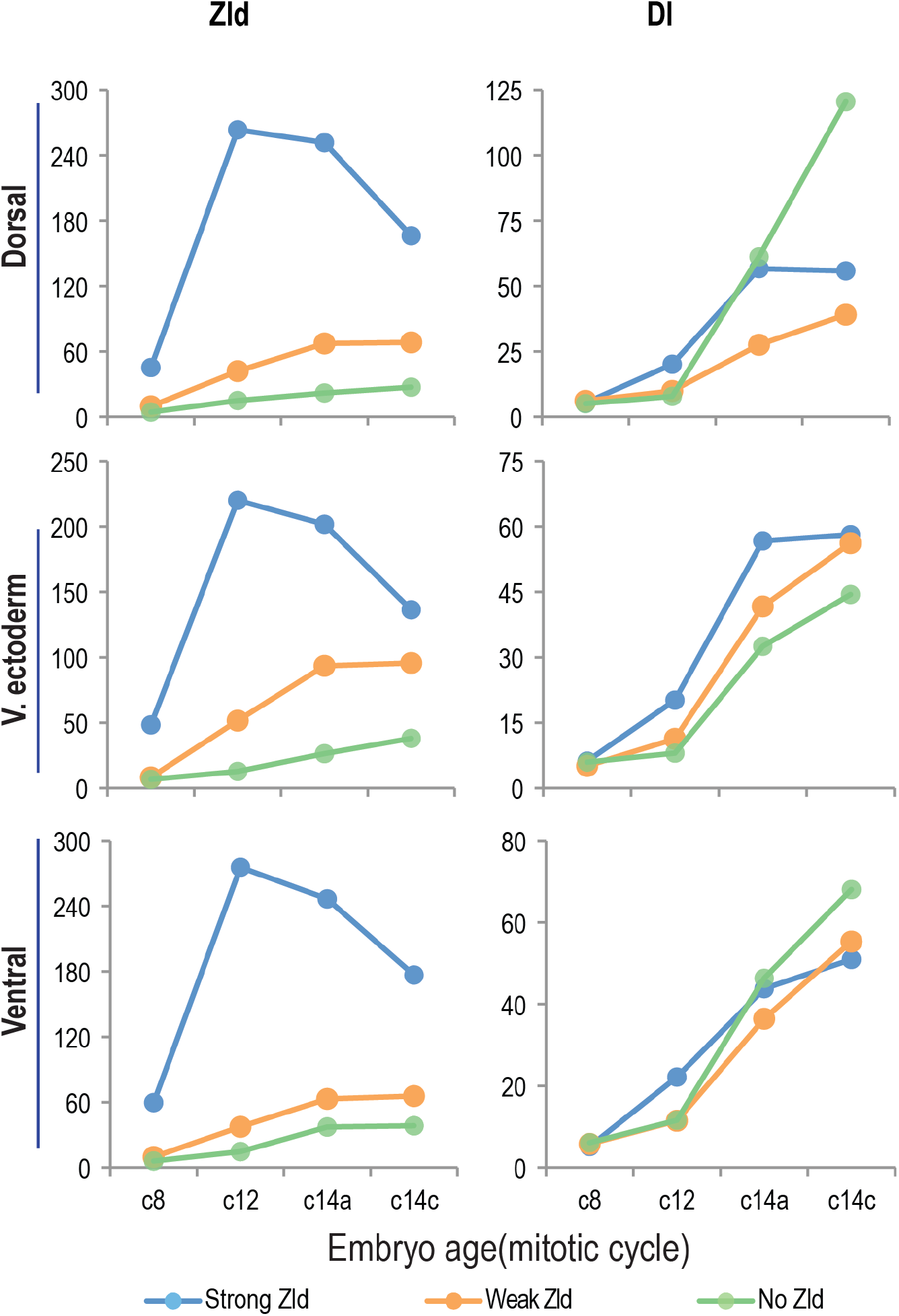
Trends of Dl binding at Dl peaks that are associated with different levels of Zld. The Dl peaks associated each of the three different classes of dorsal - ventral genes (Fig. 3A) were divided into three groups based on Zld binding at cycle 12 (Fig. 3B).

**Fig. S4.**
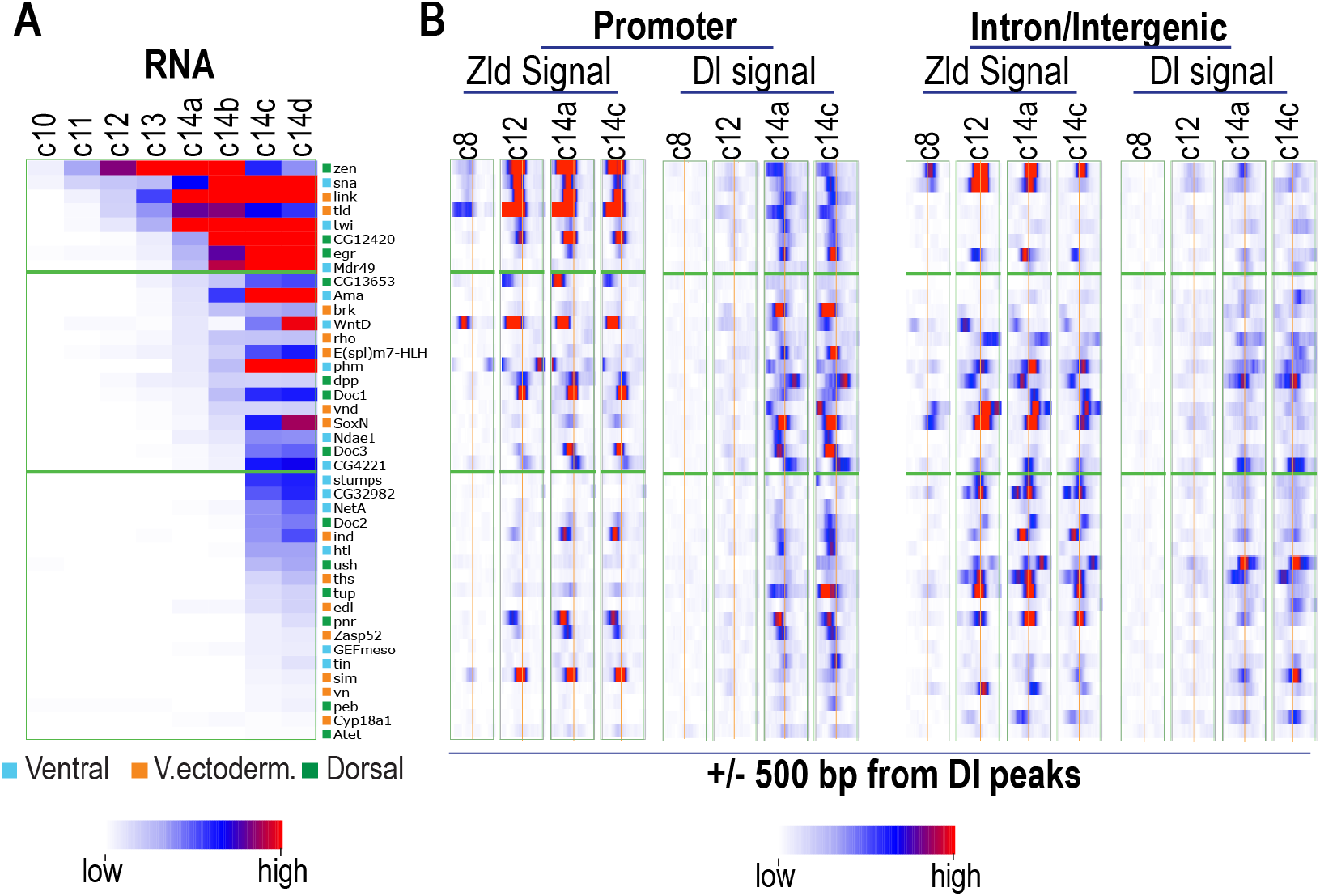
Zld binding at promoter and promoter distal regions and onset of dorsal ventral gene expression during the zygotic genome activation. A. Temporal dynamics of the transcript levels of the dorsal ventral genes based on single embryo RNA-seq analysis between c10-14 (Lott et al., 2011). The dorsal ventral genes are as described in Fig.3A; and only those genes that are expressed zygotically but not maternally are shown. Their expression patterns are as indicated on the right. The genes are sorted based on the onset time of their expression. B. Heatmap of ChIP-seq signal of Dl and Zld around Dl peaks (+/− 500bp) that are located at the promoter or promoter distal intergenic and intron regions (+/− 15 kb from transcription start site). For genes with multiple TSSs that are bound by RNA pol II, the one with the highest Dl binding is shown. Similarly, for genes with multiple promoter distal peaks, only one with the highest Dl signal is shown.

**Fig. S5.**
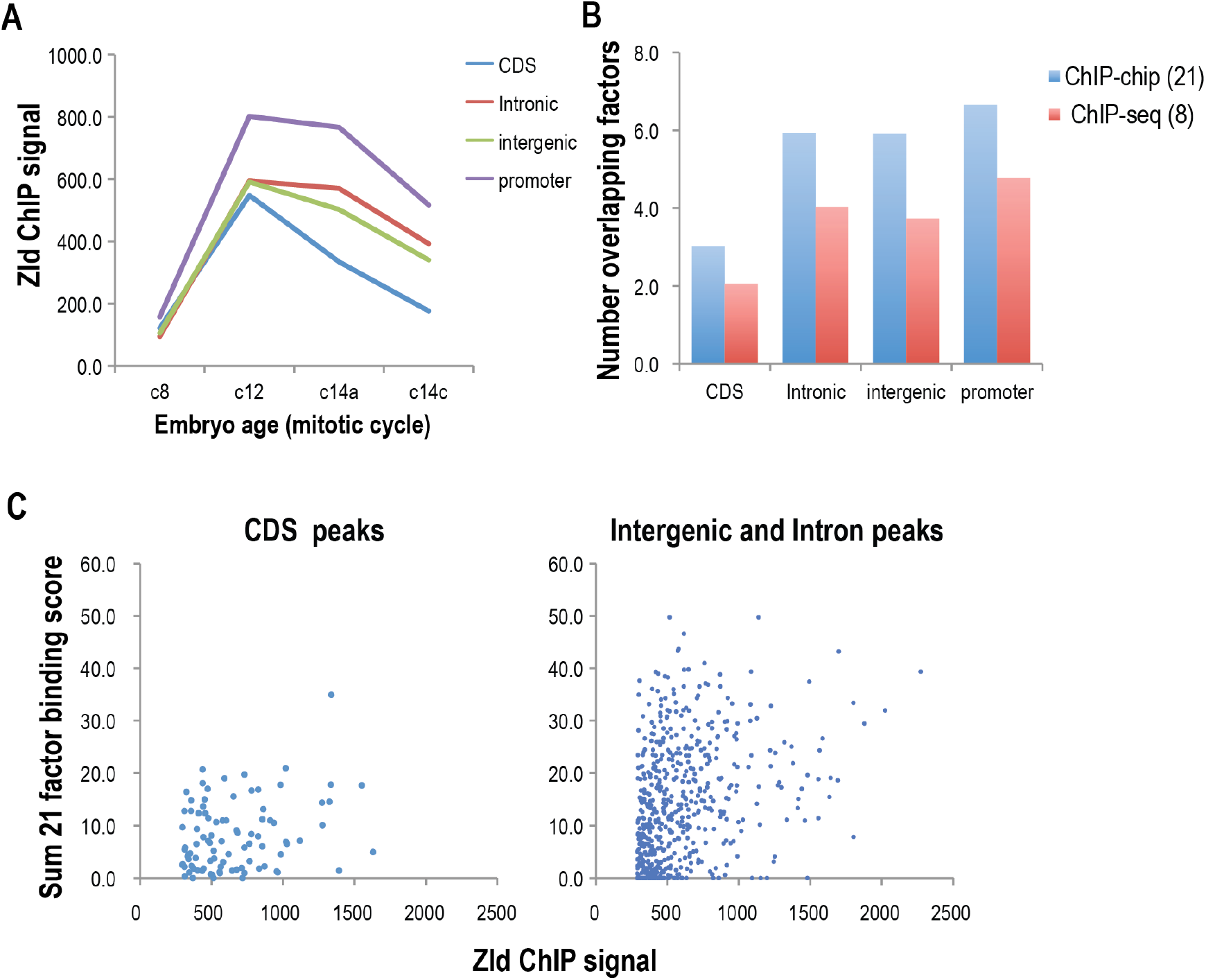
Zld peaks in coding sequences. A. Dynamics of ChIP signal of Zld peaks (top 1000 based on signal at cycle 12) located in different types of genomic regions. B. overlap of Zld peaks with transcription factors in the early embryo transcription network. Two datasets were used in the analysis: one of which included the 21 factors that have been studied by ChIP-chip (MacArthur et al., 2009), and the other included the 8 factors that have been studied by ChIP-seq including 6 A-P factors (Bradley et al., 2010), Dl (this study), and twi(He et al., 2011). C. Distribution of the sum of enrichment scores of the 21 factors for each Zld peak in the CDS. D. Distribution of the sum of enrichment scores of the 21 factors for each Zld peak in the intergenic and intron regions.

**Fig. S6.**
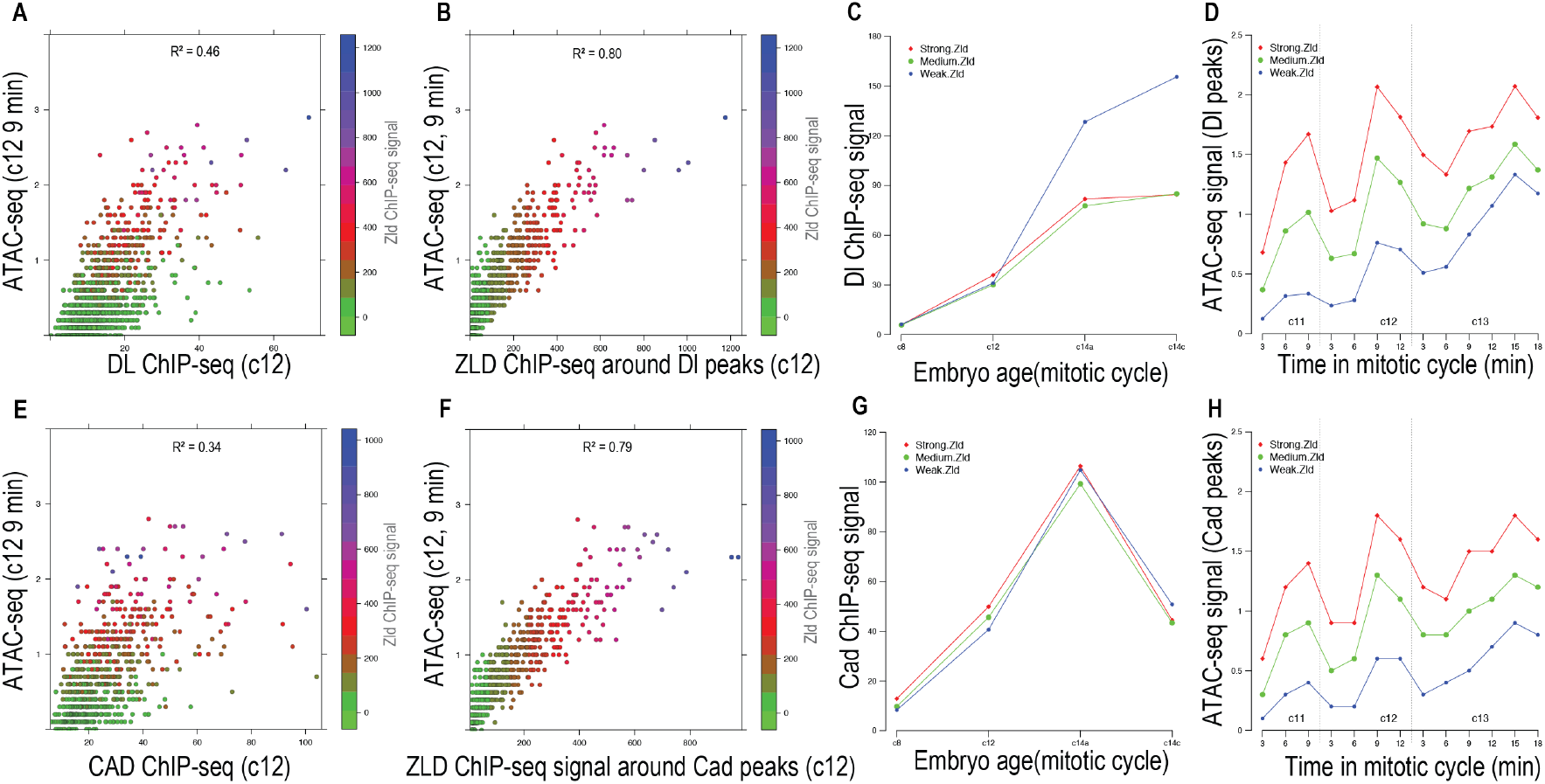
Zld binding correlates with stronger chromatin accessibility signal than Cad and Dl. A. Scatter plot of Dl ChIP signal around Dl peaks vs the ATAC-seq signal detected at 9 min in cycle 12. B. Scatter plot of Zld ChIP signal around the Dl peaks vs the ATAC-seq signal detected at 9 min in cycle 12. C. The early Dl peaks described in Fig.2 were divided into three groups according to Zld signal at cycle 12. The plot shows the trend of the average Dl ChIP signal in each of the three groups. D. The average level at each development time point and trends of the ATAC-seq signals of the three groups of peaks. E,F,G,H are plots similar to A,B,C,D, except they represent Cad peaks. The ATAC-seq data used was from (Blythe and Wieschaus, 2016)

